# Early atrial remodelling drives arrhythmia in Fabry Disease

**DOI:** 10.1101/2024.08.13.607853

**Authors:** Ashwin Roy, Christopher O’Shea, Albert Dasi I Martinez, Leena Patel, Max Cumberland, Daniel Nieves, Hansel Canagarajah, Sophie Thompson, Amar Azad, Anna Price, Caitlin Hall, Amor Mia Alvior, Phalguni Rath, Ben Davies, Blanca Rodriguez, Andrew P Holmes, Davor Pavlovic, Jonathan N Townend, Tarekegn Geberhiwot, Katja Gehmlich, Richard P Steeds

## Abstract

**Background:** Fabry disease (FD) is an X-linked lysosomal storage disorder caused by α-galactosidase A (α-Gal A) deficiency, resulting in multi-organ accumulation of sphingolipid, namely globotriaosylceramide (Gb3). This triggers ventricular myocardial hypertrophy, fibrosis, and inflammation, driving arrhythmia and sudden death, a common cause of FD mortality. Atrial fibrillation (AF) is common in FD, yet the cellular mechanisms accounting for this are unknown. To address this, we conducted electrocardiography (ECG) analysis from a large cohort of adults with FD at varying stages of cardiomyopathy. Cellular contractile and electrophysiological function were examined in an atrial FD model, developed using gene-edited atrial cardiomyocytes and imputed into *in-silico* atrial models to provide insight into arrhythmia mechanisms.

**Methods:** In 115 adults with FD, ECG P-wave characteristics were compared with non-FD controls. Induced pluripotent stem cells (iPSCs) were genome-edited using CRISPR-Cas9 to introduce the *GLA* p. *N215S* variant and differentiated into atrial cardiomyocytes (iPSC-CMs). Contraction, calcium handling and electrophysiology experiments were conducted to explore proarrhythmic mechanisms. A bi-atrial *in-silico* model was developed with the cellular changes induced by *GLA* p. *N215S* iPSC-CMs.

**Results:** ECG analysis demonstrated P-wave duration and PQ interval shortening in FD adults before onset of cardiomyopathy on imaging and biochemical criteria. FD patients exhibited a higher incidence of premature atrial contractions and increased risk of developing AF. In our cellular model, *GLA* p. *N215S* iPSC-CMs were deficient in α-Gal A and exhibited Gb3 accumulation. Atrial *GLA* p. *N215S* iPSC-CMs demonstrated a more positive diastolic membrane potential, faster action potential upstroke velocity, greater burden of delayed afterdepolarizations, greater contraction force, slower beat rate and dysfunction in calcium handling compared to wildtype iPSC-CMs. Inputting these changes into the *in-silico* model resulted in similar P-wave morphology changes to those seen in early FD cardiomyopathy and increased the action potential duration (APD) restitution slope, causing APD alternans and inducing AF.

**Conclusions:** These findings enhance our understanding of atrial myopathy in FD by providing novel insights into underpinning mechanisms for atrial arrhythmia and a rationale for early P-wave changes. These may be targeted in future research to develop therapeutic strategies to reduce the arrhythmic burden in FD and other atrial cardiomyopathies.

## INTRODUCTION

Fabry disease (FD) is a lysosomal storage disorder due to pathogenic variants in the galactosidase-α (*GLA*) gene resulting in the deficiency of α-galactosidase A (α-GAL A) (1). This results in the accumulation of glycosphingolipids, in particular globotriaosylceramide (Gb3), in lysosomes of multiple organs (2), with cardiovascular involvement the most common cause of morbidity and mortality (3). Glycosphingolipid accumulation takes place in all cardiac cell types, triggering left ventricular hypertrophy (LVH), myocardial fibrosis and inflammation, and arrhythmia. Global prevalence is estimated to range from 1:40,000 to 1:170,000 live births (4). Symptoms such as palpitations are common and occur in up to 50% of women and 75% of men. The incidence of atrial fibrillation (AF) is high at over 12% (5). Multiple structural changes are thought to contribute to the causation of AF including passive left atrial (LA) dilatation due to elevated left ventricular end-diastolic pressure (LVEDP), as well as direct atrial Gb3 accumulation resulting in atrial myopathy (6). In all populations, AF is associated with greater morbidity, including stroke, heart failure, and cardiovascular mortality (7). The risks from AF are high in those with FD, with stroke prevalence reported up to 6.9% in males and 4.3% in females (8). Furthermore, heart failure symptoms are reported in up to 25% of FD patients in AF, in large cohort studies (9).

Electrical changes can be detected early in FD on 12-lead electrocardiography (ECG) (10). Preliminary evidence from small cohorts of FD patients without LVH, suggest shortening of the P-wave duration and QRS-width, indicative of accelerated depolarization. Increased repolarization times have also been demonstrated in the same cohort manifesting as QTc prolongation (10–12). However, detailed ECG analysis measured at varying disease stages in larger cohorts of FD patients is currently lacking.

The cellular mechanisms underpinning pro-arrhythmic atrial electrical remodelling in FD are currently unknown. Enhancing our mechanistic understanding of atrial involvement in FD could identify early therapeutic targets, allow for robust risk-stratification for stroke, and may trigger initiation of disease-specific therapy to reduce the burden of cardiac complications.

We conducted signal averaged P-wave analysis from ECGs acquired in adults with FD and correlated findings with an atrial model of FD using genome edited induced pluripotent stem cell-derived atrial cardiomyocytes (iPSC-CMs). To provide further insights, we developed *in-silico* atrial models with clinical and cellular data imputed to understand how they may provoke arrhythmia.

## MATERIALS AND METHODS

### 12-lead electrocardiography (ECG)

Routinely collected 12-lead ECGs were acquired from115 adults with FD attending the centre for rare disease at the Queen Elizabeth Hospital, Birmingham between July 2014 and November 2023. The most recent ECGs were obtained from patients in sinus rhythm, to include those who subsequently developed AF. Patients were scored for cardiac disease stage using a combination of biochemical, imaging, and electrocardiographic parameters as previously described (13). Two Fabry experts (AR and RPS) reviewed the staging of patients, with discrepancies resolved by consensus. ECGs were also acquired from 40 age/sex-matched healthy non-FD controls for comparison.

ECGs were independently analysed by two experienced readers (CO and HC). Firstly, the presence of premature atrial complexes (PACs) was quantified. Secondly, 10 seconds of the lead II signal from the ECG portable document formats (PDFs) were digitised using an in-house developed algorithm and analyzed as previously described (14). Briefly, all R-waves were identified and averaged to produce the averaged ECG complex. The two observers then individually identified P-wave start and end, QRS start and end, and T-wave start and end. The isoelectric line was defined from start to end of the P-wave, and duration and amplitude parameters automatically measured.

### Transthoracic Echocardiography (TTE)

TTE data were routinely collected by an accredited sonographer (AMA) using ie33 and EPIC ultrasound systems (Phillips) according to the British Society for Echocardiography minimum dataset (15). Biplane left atrial (LA) volume was collected from the TTE report where available.

### Haematology and Biochemistry

Data for haematology and biochemistry tests performed at the nearest date to ECG acquirement were additionally extracted and included high-sensitivity (HS) troponin I, and N-terminal pro B-type natriuretic peptide (NTpro-BNP). NTpro-BNP was measured by sandwich immunoassay with magnetic particle separation and chemiluminescent detection on an E170 analyser (Roche Diagnostics, Burgess Hill, United Kingdom)

### Generation of genome-edited iPSCs

iPSCs were engineered with the pathogenic variant via CRISPR-Cas9 mediated genome-engineering to introduce the *GLA* p. *N215S* variant into male KOLF2 iPSC line as detailed in (**Supplementary Methods 1.0**) *GLA* p. *N215S* gives rise to a predominantly cardiac phenotype in FD, without overt effects in other organs (16). It is the most prevalent mutation in the United Kingdom (17). As wildtype (WT) controls, cells which were not carrying the *GLA* variant but had undergone the same genome editing procedure were used. Three independent clones were established, checked for pluripotency and correct karyotype (**Supplementary Figures 1-3**).

### Generation of atrial iPSC-derived cardiomyocytes

*GLA* p. *N215S* iPSCs were differentiated into atrial iPSC-CMs using an established protocol (18) (**Supplementary Figure 4**). Under the same conditions, WT iPSCs were differentiated into atrial iPSC-CMs. For all investigations, a minimum of three independent batches of WT and *GLA* p. *N215S* iPSC-CMs were utilized.

### Confirmation of model

The model was confirmed and validated by Western blotting for α-GAL A deficiency in *GLA* p. *N215S* iPSC-CMs, immunofluorescence for Gb3 over-accumulation in *GLA* p. *N215S* iPSC-CMs and quantitative polymerase chain reaction (qPCR) for expression of atrial and ventricular markers. Detailed methods are described in (**Supplementary Methods 2.0, 3.0 and 4.0**).

### Assessment of electrophysiological properties: Patch clamping

Stimulated and spontaneous APs were recorded from iPSC-CMs using a manual current clamp configuration. iPSC-CMs were seeded onto Geltrex (Thermo Fisher Scientific A1413201) coated coverslips 30 days after the initiation of differentiation. APs were recorded at 37 °C using an internal solution containing (in mM): KCl 135, NaCl 10, MgATP 5, HEPES 10, EGTA 0.1, pH 7.2 (adjusted using KOH). Micropipette resistances were between 3-4 MΩ. The extracellular solution contained (in mM): NaCl 145, KCl 5.4, MgSO_4_.7H_2_O 0.83, NaH_2_PO_4_ 0.33, HEPES 5, Glucose 11, CaCl_2_ 1.8, pH 7.4 (NaOH). A 60 second period after breakthrough allowed cell stabilization. Spontaneous APs were recorded for 60 seconds prior to stimulation. A continuous hyperpolarizing current was then applied to the cells to hold the diastolic membrane potential at – 75 mV prior to recording of stimulated APs (19, 20). Only those cells that required a hyperpolarizing current of less than 150 pA were used (19). APs were stimulated using a 1 nA, 2 ms current injection at a frequency of 1 Hz (20).10 successive spontaneous and stimulated APs were analyzed from each cell using custom algorithms developed in MatLab.

Delayed afterdepolarizations (DADs) may develop after the repolarization phase of an action potential, are triggered by dysregulated Ca^2+^ homeostasis, and are thought to be pro-arhythmic in nature. These were visualized and quantified during analysis. DADs were defined as a sustained spontaneous depolarisation of greater than or equal to 10mV taking place following terminal repolarization, during the diastolic interval.

### Assessment of contractile properties

Contractile properties were assessed using a GoPro Inc. camera at 20x magnification. Video recordings were obtained in multiple areas of the same well and for multiple wells of beating iPSC-CM monolayers. These were 20 second videos recorded at room temperature in a quiet setting with no other activity in the room of recording to ensure minimal interference. Videos were converted to TIFF format using DaVinci Resolve (Blackmagic Design). A TIFF-stack and single sub-stack file was then created using a script run in ImageJ. The sub-stack file was then run using a MUSCLEMOTION (21) script on ImageJ (add-on Macro); a validated tool for quantitative analysis of cardiac contraction by determining changes in pixel intensity between image frames. Outputs expressed were measures of movement during contraction and relaxation. Three 20-second videos were taken in each well with six wells for each batch The MUSCLEMOTION script generates a graphical representation of contraction and relaxation; expressed outputs are outlined below:

- Contraction amplitude – Peak contraction force from beginning of wave.
- Peak amplitude – Peak contraction force from 0.
- Relaxation time – Interval between peak contraction and beginning of following contraction.
- Time to peak - Interval between beginning of contraction and peak contraction.
- peak to-peak - Interval between each contraction.

### Assessment of calcium transients: Optical mapping

At day 17 of the differentiation protocol, atrial iPSC-CMs were split into dishes to form a confluent beating monolayer of atrial iPSC-CMs. Between days 23 and 27, calcium optical mapping was performed using intracellular calcium dye Fura-2-AM (Invitrogen). Cells were incubated with 5 µM Fura-2-AM in 1 mL RPMI B27 + Insulin (Thermo Fisher Scientific 17504044) at 37°C for 20 minutes, followed by a further 20 minutes in Tyrode’s solution (Connection in mM: NaCl 129, KCl 5.4, HEPES 10, MgCl_2_ 48, CaCl_2_ 1.8, and D-Glucose 9.99, Ph 7.44-7.48).

Dye loaded cells were excited at 380 nm, and emission imaged at 10x magnification through 510/40 nm bandpass filter. Images were collected at 1.7 ms exposure time (588 Hz) using an Evolve delta 512x512 EMCCD camera. Binning was set to 10, giving a final resolution of 51x51 pixels, pixel size = 8 µm. Images were collected for 10 to 30 seconds using WinFluor (University of Strathclyde, UK), and converted to .MAT files for analysis with ElectroMap (22). Images were pre-conditioned with a 3x3 gaussian spatial filter. Time to peak was measured from 10 to 90% calcium intrusion before peak. Calcium transient duration (CTD) was measured to 30, 50 and 80% extrusion from both maximum upstroke time (the maximum positive differential) and peak amplitude.

### Computer modelling and simulation of human atrial electrophysiology

The methodology employed for modelling and simulation is detailed in **Supplementary Methods 5.0** and briefly summarized here. *In-silico* populations of atrial cardiomyocyte models were developed to replicate and provide a mechanistic explanation to any identified differences between *GLA* p. *N215S* and WT atrial iPSC-CMs. The single-cell properties of cardiomyocyte models capturing any electrophysiological changes induced by *GLA* p. *N215S* iPSC-CMs were used to describe the electrical activity of a human bi-atrial model. This bi-atrial anatomy presented the characteristic LA (and equivalent right atrial) volume of patients showing PACs on the ECG. PACs were also replicated in-silico, by assessing changes in the P-wave morphology after applying ectopic stimuli in different locations of the human atrial model (i.e., crista terminalis and pulmonary veins). The comparison of the P-wave morphology between *in-silico* and clinical PACs was used to estimate the initiation side of abnormal electrical impulses. Subsequently, the arrhythmogenicity of the electrophysiological substrate induced by FD was assessed by applying burst stimulation on the ectopic sides identified as generators PACs. The three-dimensional monodomain equation of the transmembrane voltage and all ECG calculations were solved using the high-performance open-source MonoAlg3D (23).

### Statistical Analysis

Where two variables were being compared, normality was assessed using the Shapiro-Wilk test. If the data was normally distributed and groups had similar standard deviation (SD), unpaired t-tests were performed. If the data was normally distributed with unequal SDs, unpaired t-tests with Welch’s correction were performed. Where data was not normally distributed, the Mann-Whitney test was used. Where more than two variables were being compared, an Ordinary one-way ANOVA was performed with Turkey’s multiple comparisons test for normally distributed data. For non-normally distributed data where more than two variables were compared, the Kruskal-Wallis test was performed. For categorical and contingency data, Fishers exact test was performed. Continuous variables are reported as mean ± SD where approximately normally distributed, with non-normal variables reported as median (interquartile range; IQR) unless stated otherwise. Data was analysed and presented in GraphPad Prism version 10.0.0 with p<0.05 deemed to be indicative of statistical significance throughout.

## STATEMENT OF ETHICS

The use of 12-lead ECGs in the FD cohort was approved by West Midlands – South Birmingham Research Ethics Committee (23/WM/0180 IRAS 325613). The studies were conducted in accordance with the local legislation and institutional requirements. The Ethics Committee / institutional review board waived the requirement of written informed consent for participation from the participants or participant’s legal guardians / next of kin because data were acquired from a research database using routinely collected clinical data for the purpose of research. The use of 12-lead ECGs in healthy controls was approved by the West Midlands Solihull Research Ethics Committee (17/WM/0048) and approved by the Health Research Council. All healthy controls gave informed consent to take part in accordance with the principles set out in the Declaration of Helsinki.

iPSCs were engineered with the pathogenic *GLA* p. *N215S* variant with the support of the Genome Engineering Core Facility at the Wellcome Centre for Human Genetics, University of Oxford. The variant was introduced into male iPSCs (KOLF2 line). KOLF2 are Human Induced Pluripotent Stem Cell Initiative lines from a consortium at the Sanger Institute. They do not fall under the Human Tissue Act (2004) as the volunteer has given prior consent under an open access agreement.

## RESULTS

### Early atrial changes on ECGs of FD patients

The cohort demographics and clinical characteristics are described in (**Table 1**). The patients were predominantly female, of middle age and white Caucasian which reflects our clinical patient population. 43 (37%) had a classical FD mutation and 72 (63%) had the *GLA* p. *N215S* non-classical cardiac variant. At the time of ECG acquisition, 35 (30%) patients were on enzyme replacement therapy (ERT) and 26 (23%) on oral chaperone therapy (OCT). AF was present in 16 (14%) and therefore the ECG analyzed was that recorded when they were last in sinus rhythm. Age, male predominance, body-mass index, systolic BP and LA volume increased with cardiac disease stage. There was also a trend to worsening of cardiac, renal and AF-associated biomarkers with elevations in NTpro-BNP, HS troponin I and serum creatinine with cardiac disease stage.

**Table 1:**
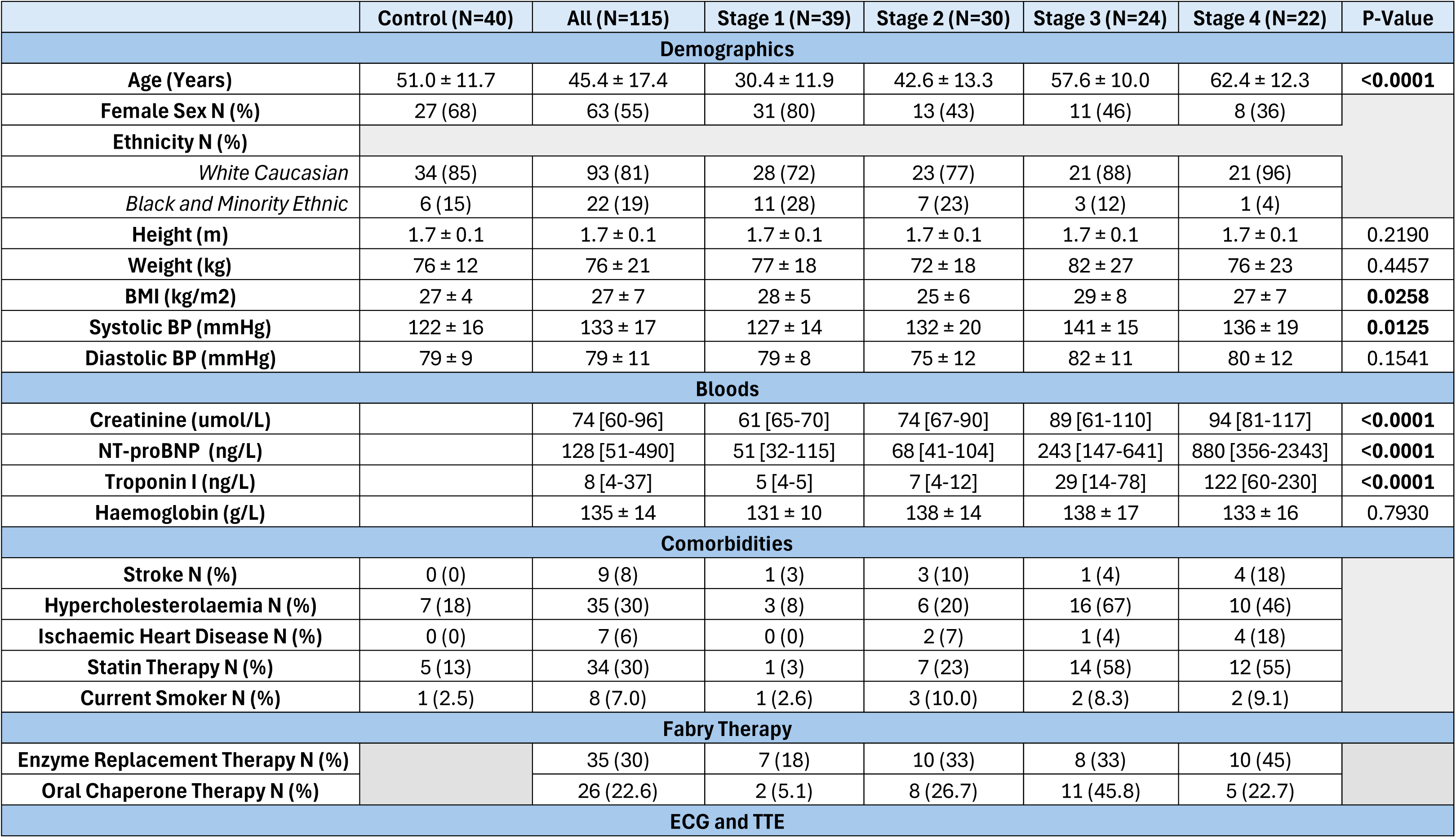
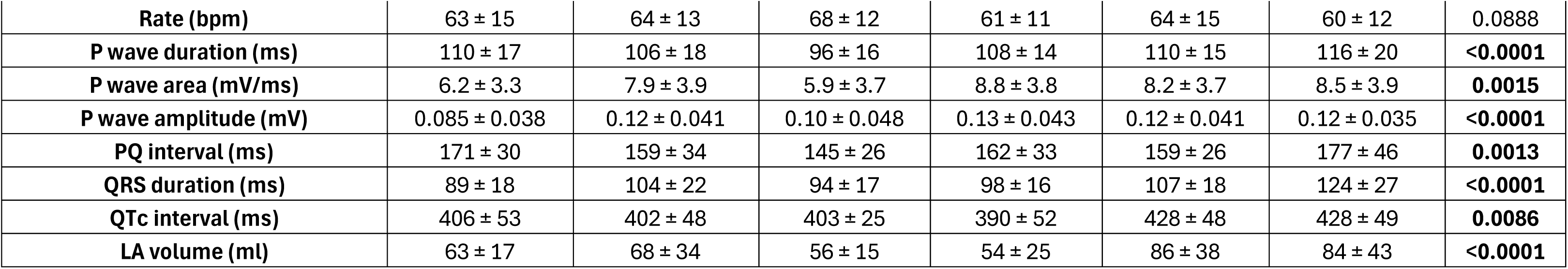
Cohort Characteristics of FD and Healthy control cohorts, total and according to cardiac disease stage. BMI: body mass index, BP: Blood pressure, LA: left atrium, ECG: electrocardiography, TTE: transthoracic echocardiography, NT-proBNP: N-terminal pro brain natriuretic peptide. Data presented as N (%) or mean ± SD. P-values derived from ordinary one-way ANOVA performed for normally distributed data. Kruskal-Wallis test performed for non-normally distributed data.

|  | Control (N=40) | All (N=115) | Stage 1 (N=39) | Stage 2 (N=30) | Stage 3 (N=24) | Stage 4 (N=22) | P-Value |
| --- | --- | --- | --- | --- | --- | --- | --- |
| Demographics |  |  |  |  |  |  |  |
| Age (Years) | 51.0 ± 11.7 | 45.4 ± 17.4 | 30.4 ± 11.9 | 42.6 ± 13.3 | 57.6 ± 10.0 | 62.4 ± 12.3 | <0.0001 |
| Female Sex N (%) | 27 (68) | 63 (55) | 31 (80) | 13 (43) | 11 (46) | 8 (36) |  |
| Ethnicity N (%) |  |  |  |  |  |  |  |
| White Caucasian | 34 (85) | 93 (81) | 28 (72) | 23 (77) | 21 (88) | 21 (96) |  |
| Black and Minority Ethnic | 6 (15) | 22 (19) | 11 (28) | 7 (23) | 3 (12) | 1 (4) |  |
| Height (m) | 1.7 ± 0.1 | 1.7 ± 0.1 | 1.7 ± 0.1 | 1.7 ± 0.1 | 1.7 ± 0.1 | 1.7 ± 0.1 | 0.2190 |
| Weight (kg) | 76 ± 12 | 76 ± 21 | 77 ± 18 | 72 ± 18 | 82 ± 27 | 76 ± 23 | 0.4457 |
| BMI (kg/m2) | 27 ± 4 | 27 ± 7 | 28 ± 5 | 25 ± 6 | 29 ± 8 | 27 ± 7 | 0.0258 |
| Systolic BP (mmHg) | 122 ± 16 | 133 ± 17 | 127 ± 14 | 132 ± 20 | 141 ± 15 | 136 ± 19 | 0.0125 |
| Diastolic BP (mmHg) | 79 ± 9 | 79 ± 11 | 79 ± 8 | 75 ± 12 | 82 ± 11 | 80 ± 12 | 0.1541 |
| Bloods |  |  |  |  |  |  |  |
| Creatinine (umol/L) |  | 74 [60-96] | 61 [65-70] | 74 [67-90] | 89 [61-110] | 94 [81-117] | <0.0001 |
| NT-proBNP (ng/L) |  | 128 [51-490] | 51 [32-115] | 68 [41-104] | 243 [147-641] | 880 [356-2343] | <0.0001 |
| Troponin I (ng/L) |  | 8 [4-37] | 5 [4-5] | 7 [4-12] | 29 [14-78] | 122 [60-230] | <0.0001 |
| Haemoglobin (g/L) |  | 135 ± 14 | 131 ± 10 | 138 ± 14 | 138 ± 17 | 133 ± 16 | 0.7930 |
| Comorbidities |  |  |  |  |  |  |  |
| Stroke N (%) | 0 (0) | 9 (8) | 1 (3) | 3 (10) | 1 (4) | 4 (18) |  |
| Hypercholesterolaemia N (%) | 7 (18) | 35 (30) | 3 (8) | 6 (20) | 16 (67) | 10 (46) |  |
| Ischaemic Heart Disease N (%) | 0 (0) | 7 (6) | 0 (0) | 2 (7) | 1 (4) | 4 (18) |  |
| Statin Therapy N (%) | 5 (13) | 34 (30) | 1 (3) | 7 (23) | 14 (58) | 12 (55) |  |
| Current Smoker N (%) | 1 (2.5) | 8 (7.0) | 1 (2.6) | 3 (10.0) | 2 (8.3) | 2 (9.1) |  |
| Fabry Therapy |  |  |  |  |  |  |  |
| Enzyme Replacement Therapy N (%) |  | 35 (30) | 7 (18) | 10 (33) | 8 (33) | 10 (45) |  |
| Oral Chaperone Therapy N (%) |  | 26 (22.6) | 2 (5.1) | 8 (26.7) | 11 (45.8) | 5 (22.7) |  |
| ECG and TTE |  |  |  |  |  |  |  |
| **Rate (bpm)** | 63 ± 15 | 64 ± 13 | 68 ± 12 | 61 ± 11 | 64 ± 15 | 60 ± 12 | 0.0888 |
| **P wave duration (ms)** | 110 ± 17 | 106 ± 18 | 96 ± 16 | 108 ± 14 | 110 ± 15 | 116 ± 20 | **<0.0001** |
| **P wave area (mV/ms)** | 6.2 ± 3.3 | 7.9 ± 3.9 | 5.9 ± 3.7 | 8.8 ± 3.8 | 8.2 ± 3.7 | 8.5 ± 3.9 | **0.0015** |
| **P wave amplitude (mV)** | 0.085 ± 0.038 | 0.12 ± 0.041 | 0.10 ± 0.048 | 0.13 ± 0.043 | 0.12 ± 0.041 | 0.12 ± 0.035 | **<0.0001** |
| **PQ interval (ms)** | 171 ± 30 | 159 ± 34 | 145 ± 26 | 162 ± 33 | 159 ± 26 | 177 ± 46 | **0.0013** |
| **QRS duration (ms)** | 89 ± 18 | 104 ± 22 | 94 ± 17 | 98 ± 16 | 107 ± 18 | 124 ± 27 | **<0.0001** |
| **QTc interval (ms)** | 406 ± 53 | 402 ± 48 | 403 ± 25 | 390 ± 52 | 428 ± 48 | 428 ± 49 | **0.0086** |
| **LA volume (ml)** | 63 ± 17 | 68 ± 34 | 56 ± 15 | 54 ± 25 | 86 ± 38 | 84 ± 43 | **<0.0001** |

P-wave morphology changes in non-FD controls, stage 1 FD and stage 4 FD are illustrated in (**Figure 1A**). There was a significant shortening of P-wave duration in adults with stage 1 cardiac phenotype FD compared to non-FD controls (96 ± 16 ms versus 110 ± 17 ms; p=0.0026) **(Figure 1B**). PQ interval shortening in FD stage 1 was also present when compared to non-FD controls (145 ± 27 ms versus 171 ± 30 ms; p=0.0043) (**Figure 1C**). Prolongation of P-wave duration and PQ interval were seen to be associated with worsening severity of FD cardiac phenotype (stages 1-4) (**Figures 1A-C and Table 1**). Prolongation of P-wave duration and PQ interval in the later stages of cardiac disease was associated with greater LA volume on TTE (**Figure 1D**). P-wave duration and PQ interval were also shorter in patients with stage 1 FD in the *GLA p. N215S* cohort compared to healthy controls (96 ± 18 ms versus 110 ± 17 ms; p=0.0026)

**Figure 1:**
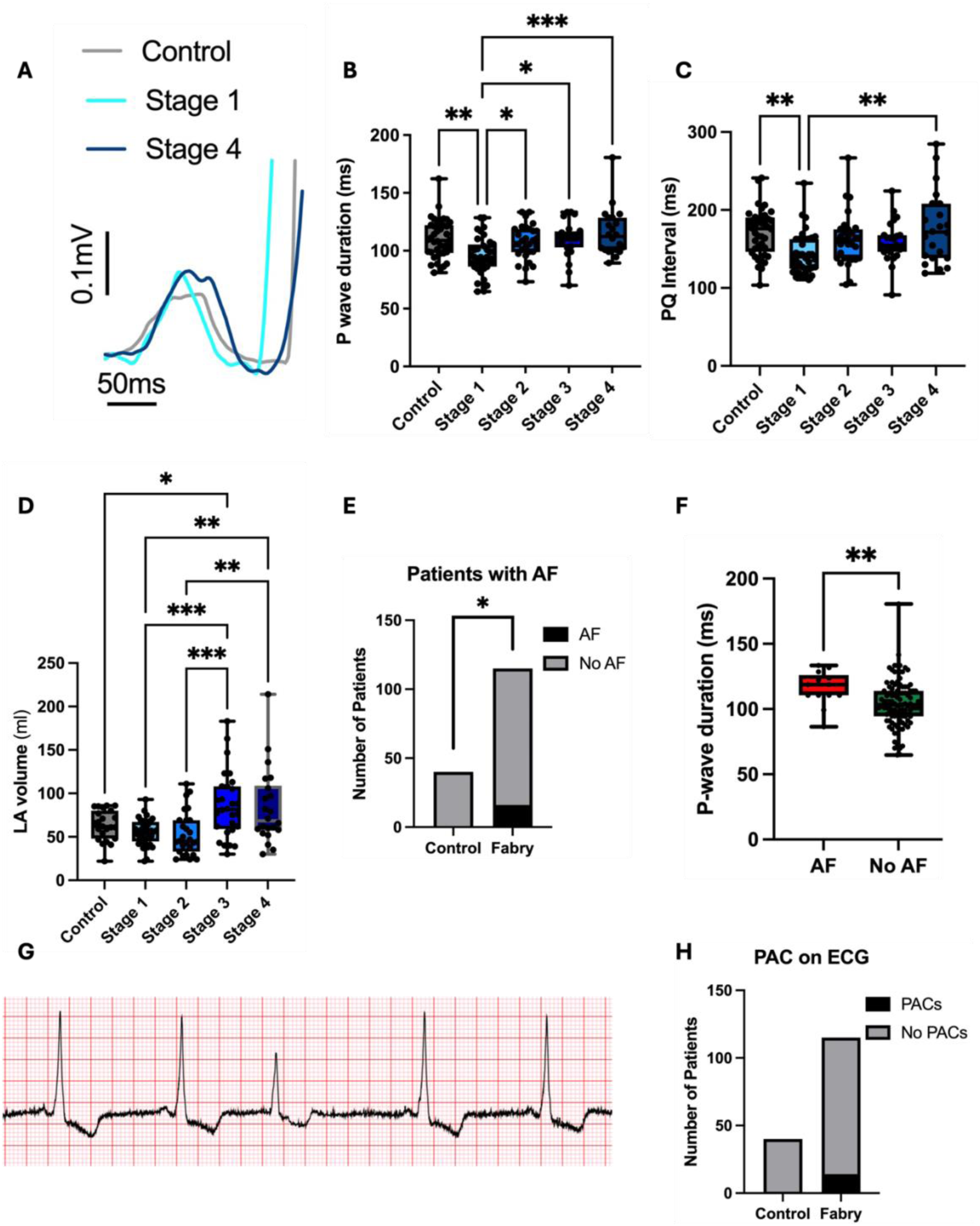
12-lead ECG analysis: (A) P-wave morphology changes on ECG (control vs stage 1 vs stage 4) (B-C) P-wave duration and PQ interval changes in controls and FD cardiomyopathy stages (D) LA volume changes in controls and FD cardiomyopathy stages on TTE. (E) Proportion of patients with AF Fabry vs control (F) P-wave duration in AF vs no AF (G) PAC on ECG of adult with FD (H) Proportion of PACs Fabry Vs control. Ordinary one-way ANOVA with Turkey’s multiple comparisons statistical test used comparing stages. Mann Whitney Test used to compare P-wave duration in AF vs no AF. Fisher’s exact test used for proportion of patients with PACs and AF. Data presented as mean ± SD. * p ≤ 0.05, ** p ≤ 0.01, *** p ≤ 0.001

More FD patients had AF compared to non-FD controls (**Figure 1E**). In the sub-analysis of patients with AF (N=16) using their most recent ECGs in sinus rhythm, a more prolonged P-wave duration was documented compared to those remaining in sinus rhythm (116 ± 13 ms versus 104 ± 18 ms; p=0.0094) (**Figure 1F**). The presence of PACs on ECG (**Figure 1G)** were observed in 14/115 adults with FD vs 0/40 for controls (**Figure 1H)**. Prevalence of PACs in cardiac disease stages were not significantly different; Stage 1 (N=4/39), stage 2 (N=1/30), stage 3 (N=5/24) and stage 4 (N=4/22).

### Successful generation of atrial FD model

Sanger sequencing confirmed the presence of *GLA* p. *N215S* at the CRISPR target site, fluorescence activated cell sorting (FACS) analysis confirmed high expression of pluripotent stem cell markers and normal chromosomal count was confirmed on karyotype analysis (**Supplementary Figures 1-3**). α-GAL A is a homodimer, consisting of two 49 kDA subunits. Denaturing western blotting confirmed reduced expression of α-GAL A in *GLA* p.*N215S* iPSC-CMs compared to WT (**Figure 2A and Supplementary Figure 5**). Significantly lower expression of α-GAL A in *GLA* p.*N215S* iPSC-CMs was also confirmed when normalized to WT (0.24 ± 0.22-fold change from WT at 1.0 ± 0.37; p=0.0368) (**Figure 2B).** When stained with a Gb3 antibody and visualised using confocal fluorescence microscopy, mature atrial *GLA* p.*N215S* iPSC-CMs displayed greater accumulation of Gb3 compared to WT (**Figure 2C-D).** On Gb3 quantification, *GLA* p.*N215S* iPSC-CMs exhibited a greater number of puncta per micron^2^ compared to WT (0.031: IQR 0.22-0.45 puncta per micron^2^ versus 0.014: IQR 0.099-0.019 puncta per micron^2^; p=0.0073) (**Figure 2E).** Markers of atrial and ventricular expression on qPCR are illustrated in (**Supplementary Figure 6A-D**). When normalized to GAPDH, atrial iPSC-CMs had greater expression of atrial marker *MYL7* (1.9: IQR 1.2-3.7-fold change in atrial iPSC-CMs versus 0.87: IQR 0.82-1.4-fold change in ventricular iPSC-CMs: p=0.0184) and a trend to increased expression of *MYH6,* with reduced expression of ventricular markers *MYL2* (0.14: IQR 0.067-0.023-fold change in atrial iPSC-CMs versus 0.62: IQR 0.39-4.1-fold change in ventricular iPSC-CMs; p=0.0021) and *MYH7* (0.31: IQR 0.11-0.91-fold change in atrial iPSC-CMs versus 0.93: IQR 0.86-1.2-fold change in ventricular iPSC-CMs; p=0.0142) compared with ventricular iPSC-CM expression. All iPSC-CMs expressed the cardiac marker troponin (*TNNT2*) on qPCR (**Supplementary Figure 6E**).

**Figure 2:**
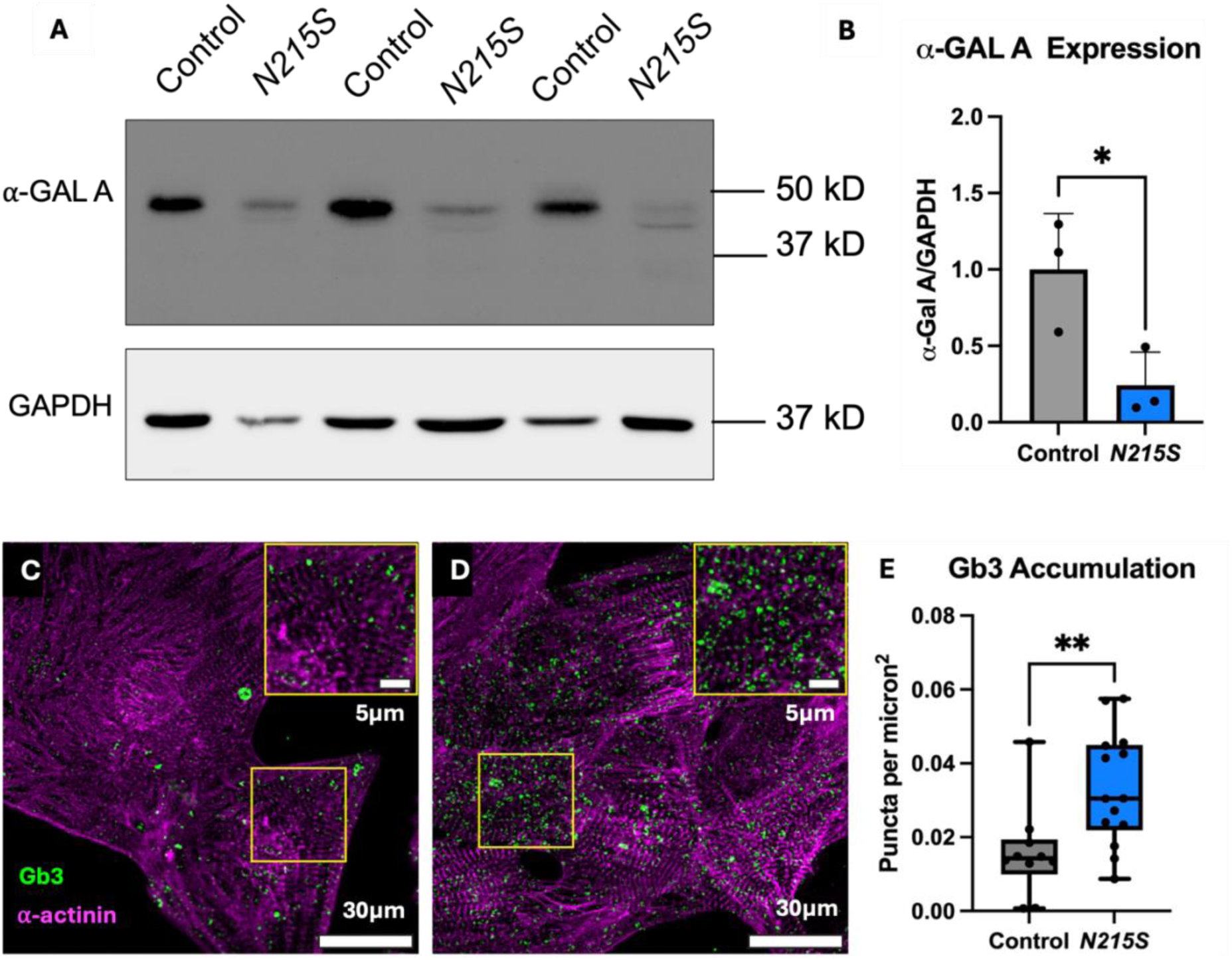
Confirmation of FD model. (A) Western blot for α-GAL A protein levels (control vs *N215S*) in iPSC-CMs. (B) Quantification of α-GAL A expression (control vs *N215S*) in iPSC-CMs. Data presented as mean ± SD (C-D) Immunofluorescence Gb3 (green) stain using confocal microscopy (A: control B: *N215S*) in atrial iPSC-CMs (α-actinin, purple) (E) Gb3 accumulation (control vs *N215S*) in atrial iPSC-CMs. Data presented as mean ± SD. Mann Whitney U statistical test used,* p=<0.05, ** p=<0.01.

### Quicker upstroke in atrial APs of GLA p. N215S IPSC-CMs identified

Findings from single cell patch clamping assessment of atrial APs are summarised in (**Figure 3**). In the first step, we assessed spontaneous action potentials. *GLA* p. *N215S* atrial iPSC-CMs demonstrated a more positive resting diastolic membrane potential (**Figure 3A**) (-49 ± 7.1 mV versus -52 ± 7.1 mV; p=0.0153). There were no significant changes in firing frequency or APD30, 50, 70 or 90.

**Figure 3:**
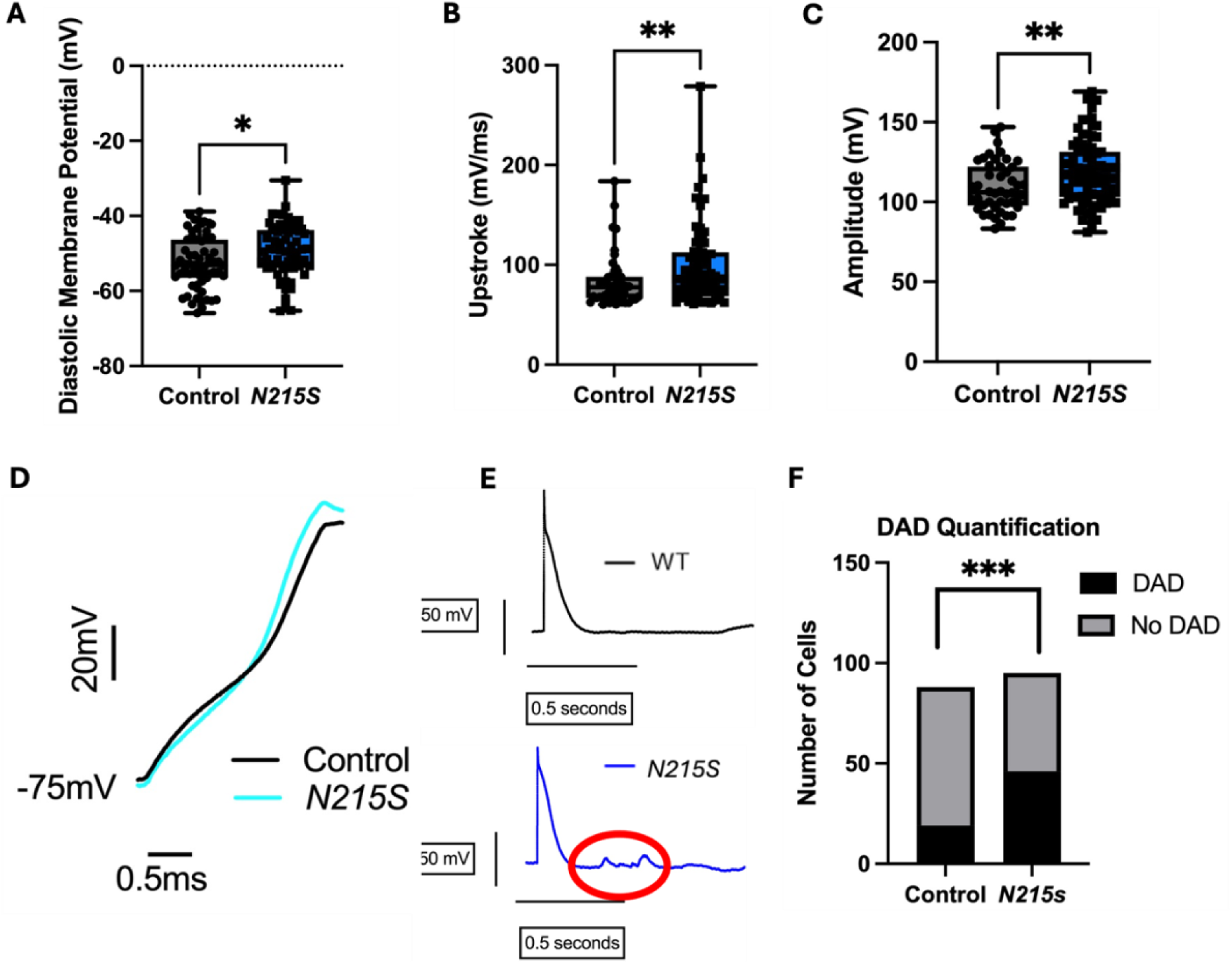
(A-C) Diastolic membrane potential, upstroke and amplitude from APs from iPSC-CMs at 1Hz (WT vs *N215S*). (D) Single cell atrial APs (WT vs *N215S*) illustrating quicker upstroke in *N215S*. (E) Stimulated atrial APs (WT vs *N215S*) illustrating DAD activity in *N215S* (circled in red) compared to no DAD activity in WT. (F) DAD quantification documenting a greater number of *N215S* iPSC-CMs with DAD activity compared to WT. Welch’s unpaired t-test statistical test used for Diastolic membrane potential. Mann Whitney U test used for Stimulated Upstroke and Amplitude. Fisher’s exact test used for comparison of number of DADs. Data presented as mean ± SD.* p ≤ 0.05, *** p ≤ 0.001, **** p ≤ 0.0001,

In the second step, we assessed stimulated APs. When stimulating cells at a held resting potential of -75 mV with 1 nA, 2 ms current injection, *GLA* p. *N215S* atrial iPSC-CMs had a significantly faster action potential upstroke velocity (87: IQR 75-113 mV/ms versus 78: IQR 67-88 mV/ms; p=0.004) and a greater action potential amplitude (118 ± 20 mV versus 110 ± 16 mV; p=0.0095) (**Figure 3B-D**). There were no significant changes in APD30, APD50, APD70 or APD90.

The presence of DADs in a *GLA* p. *N215S* atrial iPSC-CM stimulated AP are illustrated in (**Figure 3E**). Quantitative assessment of stimulated APs showed an increased numbers of cells with APs displaying DADs in *GLA* p. *N215S* atrial iPSC-CMs (N=46) compared with WT (N=19) (**Figure 3F**) and a greater number of DADs per cell, in *GLA* p. *N215S* atrial iPSC-CMs (p=0.0002).

### Greater contraction in atrial GLA p. N215S iPSC-CMs identified

Findings from MUSCLEMOTION analysis of contraction in the 2D monolayers are summarised in (**Figure 4**). Graphical representations of contraction and relaxation are illustrated for *GLA* p. *N215S* iPSC-CMs (**Figure 4A**) and WT iPSC-CMs (**Figure 4B).** Data from MUSCLEMOTION findings are expressed as normalized percentage changes of the mean of the WT. *GLA* p. *N215S* iPSC-CMs had a significantly longer contraction duration (119: IQR 103-145 % versus 98: IQR 86-115 %; p=0.0055) (**Figure 4C)** and time to peak (150: IQR 84-212 % versus 98: IQR 82-118 %; p=0.0296) (**Figure 4D),** as well as a longer peak-to-peak distance (145 ± 59 % versus 100 ± 16.9 %; p=0.0032) (**Figure 4H)** indicating a slower beat rate compared to WT. *GLA* p. *N215S* iPSC-CMs also generated a significantly higher contraction (141: IQR 103-194 % versus 101: IQR 65-129 %; p=0.0074) and peak (119: IQR 100-133 % versus 95: IQR 75-116 %; p=0.0047) amplitude (**Figure 4E** and **F**) compared to WT. There were no differences in relaxation time observed between *GLA* p. *N215S* iPSC-CMs and WT (**Figure 4G**).

**Figure 4:**
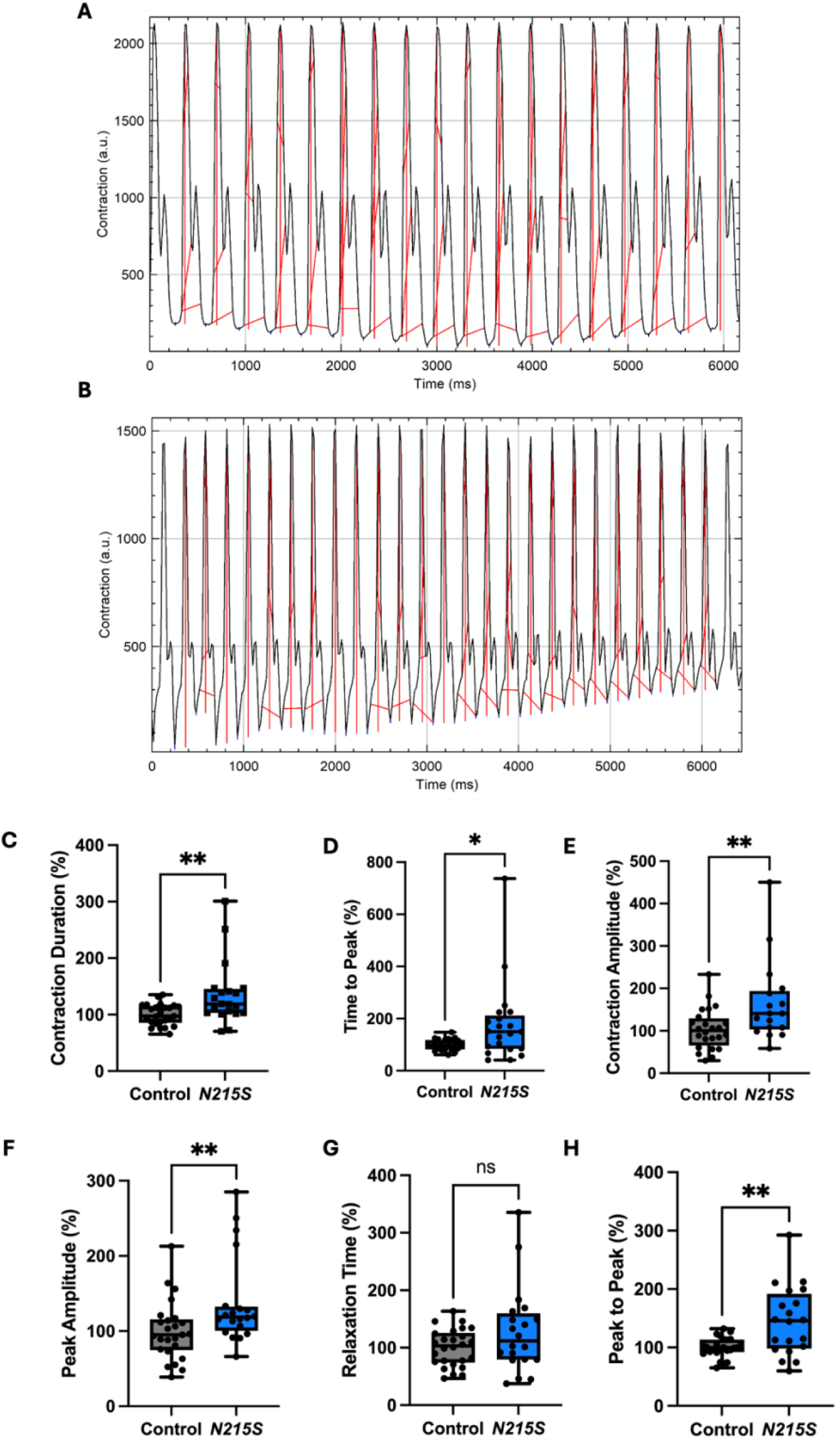
(A) Illustration of the outputs from MUSCLEMOTION in an *N215S* atrial iPSC-CM and (B) WT-iPSC-CM. (C-H) Intervals and contractility parameters from atrial iPSC-CMs (*N215S* vs WT) Values expressed as a percentage difference of the mean value for the WT of each batch. Mann Whitney U statistical test used for Contraction Duration, Time to Peak, Contraction Amplitude, Peak Amplitude and Relaxation time. Welch’s unpaired t-test used for Peak to Peak time. Data presented as mean ± standard deviation. ns not significant, * p ≤ 0.05, ** p ≤ 0.01.

### Prolonged CTDs in atrial GLA p. N215S iPSC-CMs identified

To assess calcium handling, optical mapping was performed in the cellular monolayers. CTD was prolonged to all extrusion levels in *GLA* p. *N215S* atrial iPSC-CMs as demonstrated in the polar maps of each, with a different transient morphology compared to WT controls across the monolayers (**Figure 5A-C)**: CTD30 (265 ± 35 ms versus 190 ± 37 ms; p=<0.0001), CTD50 (357 ± 37 ms versus 252 ± 54 ms; p=<0.0001) and CTD80 (587: IQR 535-647 ms versus 358: IQR 337-453 ms; p=<0.0001) (**Figures 5D-F)**. *GLA* p. *N215S* atrial iPSC-CMs also displayed a significantly slower time-to-peak (**Figure 5G)** (96 ± 26 ms versus 77 ± 18 ms; p=0.0034), and larger peak amplitude (**Figure 5H)** (16325 ± 2595 au versus 13998 ± 2894 au; p=0.0004) suggesting a greater total cycling of calcium, and a slower beating frequency (**Figure 5I)** (35: IQR 27-38 bpm versus 56: IQR 40-70 bpm; p=<0.0001) consistent with contraction analysis data.

**Figure 5:**
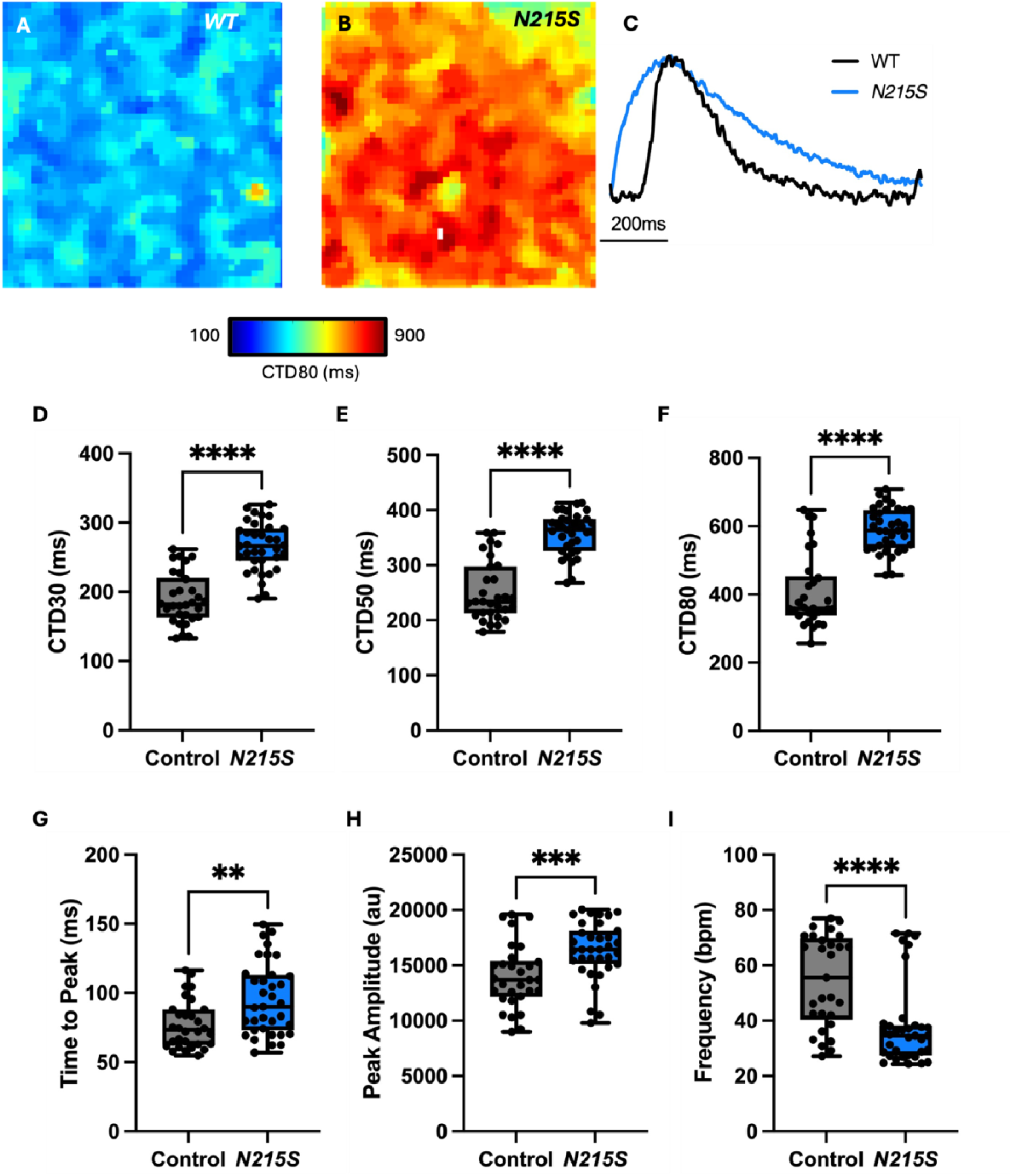
(A) WT atrial iPSC-CM CTD polar map indicating shorter CTDs (blue) (B) *N215S* atrial iPSC-CM CTD polar map indicating longer CTDs (red) (C) Atrial calcium transients (*N215S* vs WT) illustrating changes in morphology and CTDs (D-F) Atrial iPSC-CM CTD30, 50, 80 (*N215S* vs WT) (G-I) Interval and amplitude parameters (*N215S* vs WT). Mann Whitney U statistical test used for CTD80, CTD50, Time to peak, Peak Amplitude and Frequency. Welch’s Unpaired t-test used for CTD30. Data presented as mean ± standard deviation * p ≤ 0.05, ** p ≤ 0.01, *** p ≤ 0.001, **** p ≤ 0.0001.

### Action potential alternans as a mechanism of AF in patients with FD

An *in-silico* population of 222 human atrial cardiomyocyte models reproduced the faster upstroke and higher action potential amplitude observed in *GLA* p. *N215S* atrial iPSC-CMs (**Figure 6A**), as well as the slower time-to-peak, larger peak amplitude and longer duration of the calcium transient (**Supplementary Figure 7A**). Compared to 251 atrial cell models without these alterations, the calcium changes observed in *GLA*p. *N215S* atrial iPSC-CMs favoured an increase in the APD restitution slope and the appearance of APD alternans *in-silico* (**Figure 6A**). One atrial cardiomyocyte model representative of FD, with faster upstroke, higher AP amplitude and APD alternans, and one without these alterations, representative of control, were used to describe the electrophysiology of a human bi-atrial model (right and left atrial volumes of 122 mL and 85 mL, respectively). During sinus rhythm, shorter P-waves (i.e., faster atrial depolarization) were obtained in the 3D atrial anatomy representative of FD, as observed clinically in patients with FD (**Figure 6C)**

**Figure 6:**
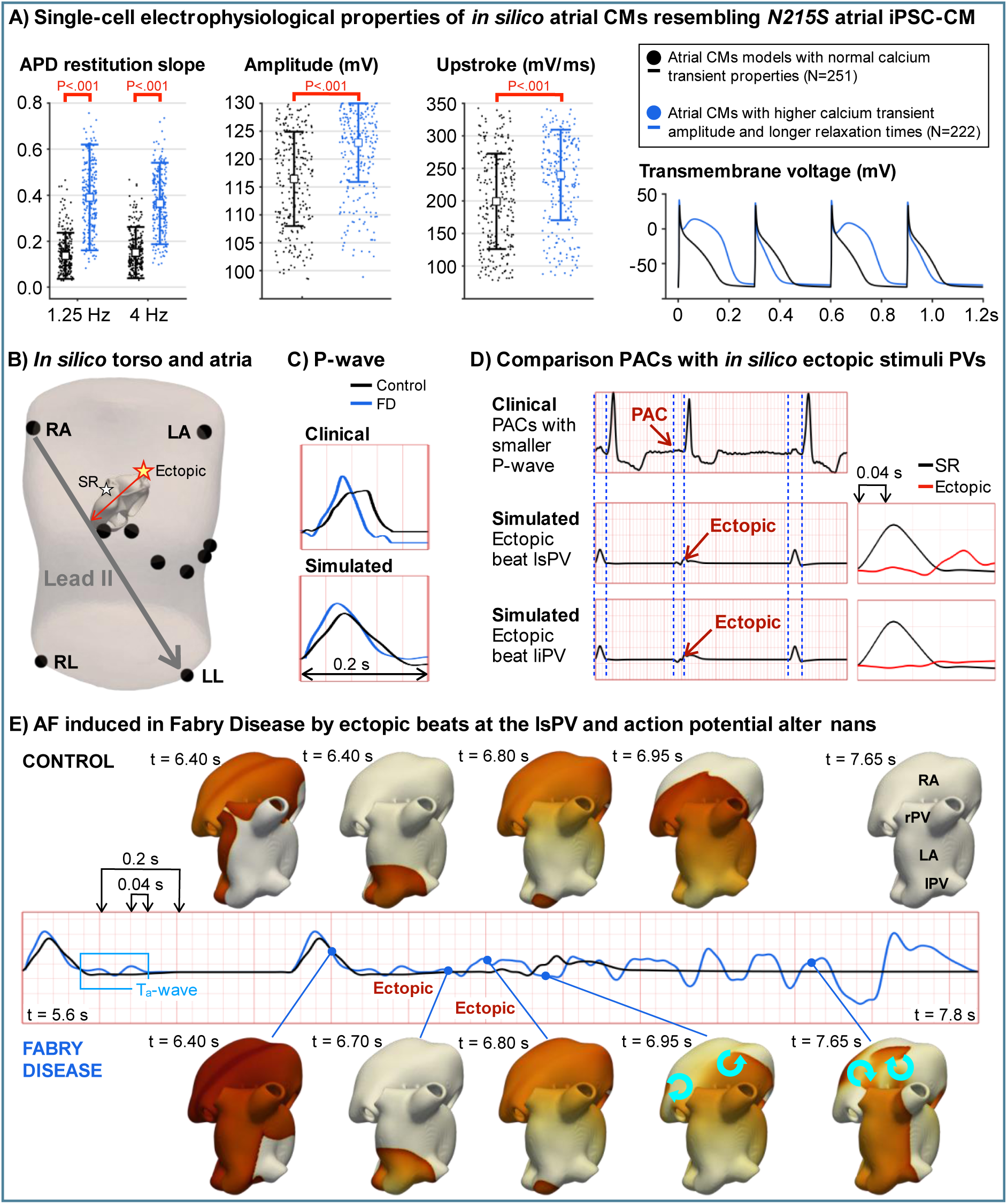
*In silico* investigation of the proarrhythmic profile of FD. (A) Action potential duration (APD) restitution slope, action potential amplitude and maximum upstroke velocity of 251 and 222 human atrial cardiomyocyte models representative of WT and *GLA* p. *N215S* atrial iPSC-CMs, respectively. Time course of the transmembrane voltage of a representative cardiomyocyte model in each subgroup. The cardiomyocyte model representative of *GLA* p. *N215S* atrial iPSC-CMs (in blue) shows action potential alternans. These atrial cardiomyocyte models, representative of FD (blue) and control (black), are used to describe the electrophysiology of a human bi-atrial model. (B) Schematic representation of the human atrial model inside a torso with the electrode location. It is illustrated the location of the sinus node (stimulation during sinus rhythm and the direction of propagation of an ectopic beat originating from the left pulmonary vein (lPV), Abbreviations: RA-LA-LL: right arm, left arm and left leg (C) Comparison of clinical and simulated P-waves, considering the representative atrial cardiomyocyte models shown in (A). Shorter P-waves are observed in the 3D anatomy with FD-induced electrophysiological changes, as observed clinically. D) Comparison between premature atrial contractions (PACs) in a patient with FD and in-silico ectopic stimuli from the lsPV. An ectopic beat originating from the lPV yields a direction of propagation perpendicular to lead II, which results in a flat P-wave, similar to that observed clinically after the PAC. (E) Consecutive snapshots of the atrial transmembrane voltage and corresponding ECG in control conditions and FD. AF is induced by two ectopic beats applied in the lsPV only when considering the electrophysiological changes caused by FD. The blue arrows on top of the atrial anatomy indicate the presence of rotors. Since the QRS is not simulated, the atrial repolarization (Ta-wave) can be seen on the ECG. Abbreviations: RA-LA: right and left atrium; rPV-lPV: right and left pulmonary veins.

This 3D atrial anatomy was used to estimate the ectopic origin of PACs in patients with FD. For this, ectopic stimuli were applied in different atrial sites of the human bi-atrial model (**Supplementary Figure 8**) and the resulting changes in the P-wave morphology were compared against PACs on ECG. The 3D atrial anatomy representative of FD was used for assessing ectopic analysis, since PACs were only observed in patients with FD (**Figure 1**). The comparison between simulated and clinical ECGs evidenced the pulmonary veins as a potential ectopic trigger of PACs in patients with FD (**Figure 6B** and **D**), in agreement with the greater number of DADs observed in *GLA* p. *N215S* atrial iPSC-CMs (**Figure 3**). Indeed, two consecutive ectopic beats applied at fast rates (e.g. 130 ms) in the left superior pulmonary vein were sufficient to induce AF in the presence of the electrophysiological changes described above (**Figure 6E**).

## DISCUSSION

In a deeply phenotyped cohort of patients with FD, staged according to degree of cardiomyopathy, we demonstrate P-wave duration and PQ interval shortening in adults with FD, even in those without an overt cardiac disease phenotype. This has not previously been demonstrated in “cardiac phenotype-negative” patients and for the first time suggests early atrial electrical remodelling secondary to Gb3 accumulation in patients with classical disease and with the *GLA* p. *N215S* variant, previously thought to be “late-onset”. P-wave duration and PQ interval shortening have been demonstrated previously in a small study of (N=30) FD patients without LVH, however these patients did not undergo multiparametric CMR and biomarker assessment to exclude early-onset cardiomyopathy due to myocardial Gb3 accumulation (10). Crista terminalis and atrioventricular (AV) nodal Gb3 accumulation may account for P-wave duration shortening via accelerated conduction (24–26). As cardiac disease stage progresses, P-wave duration and PQ interval appears to prolong, and a positive correlation is observed with LA volume on TTE. These findings suggest multiple mechanisms underlying the pathophysiology of atrial myopathy in FD. Firstly, in early disease, intrinsic electrophysiological cellular changes, presumably due to Gb3 accumulation, cause abnormalities in sodium and calcium handling that help to account for the early PQ interval and P-wave duration shortening on 12-lead ECG seen in FD patients. These may be pro-arrhythmic in nature (12). As Gb3 accumulation progresses and LA volume increases due to the passive effects of elevated LVEDP due to LVH, this structural alteration causes ‘pseudo-normalization of the P-wave duration and PQ interval, which eventually becomes prolonged, further increasing the susceptibility to the onset of AF (27). These mechanisms according to disease stage are summarised in (**Figure 7**). These are important findings as they provide insight into underpinning mechanisms and may serve as biomarkers in the assessment of treatment response.

**Figure 7:**
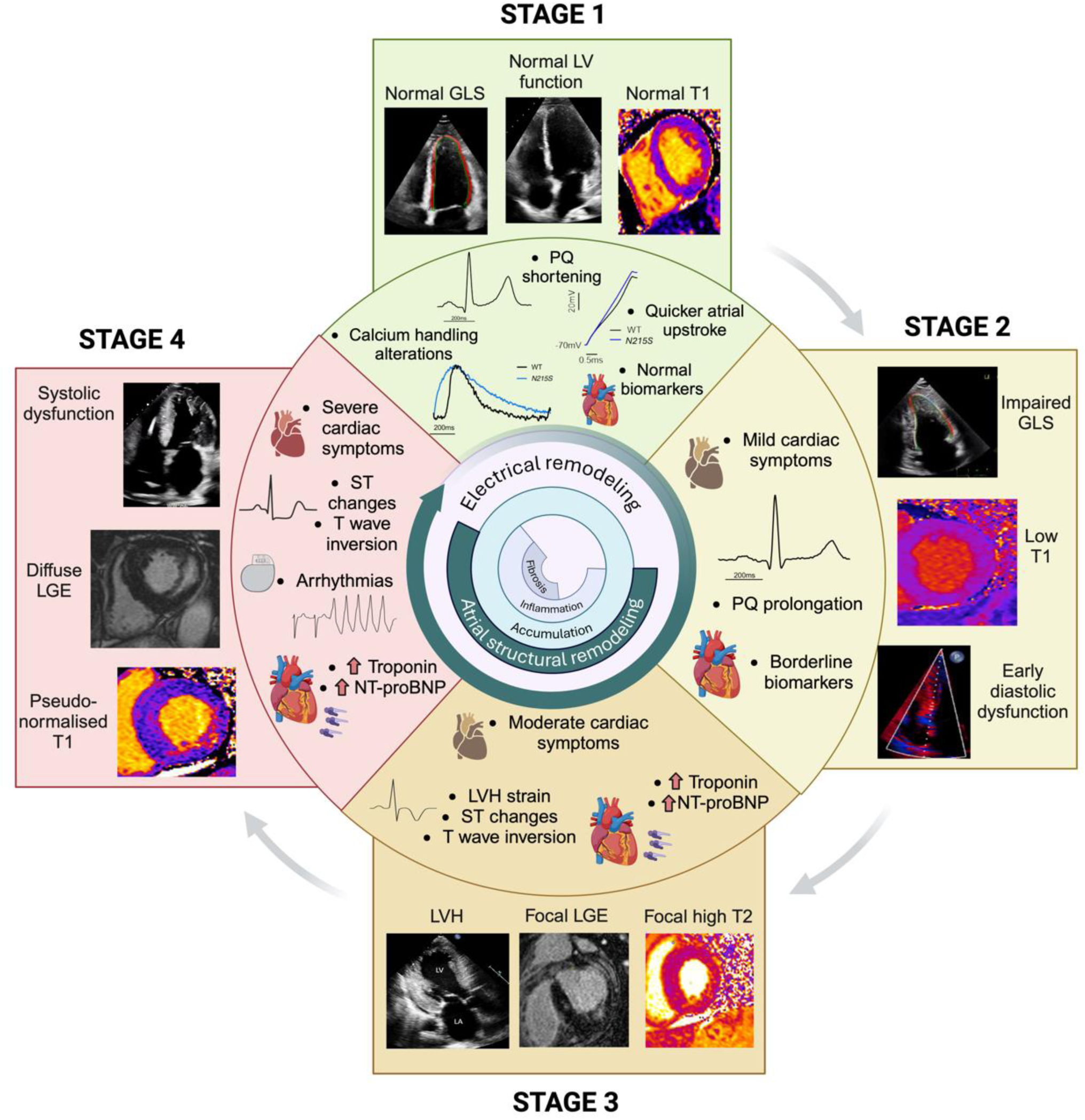
Stages of cardiomyopathy in FD. Abbreviations: GLS: global longitudinal strain, LV: left ventricular, LGE: late gadolinium enhancement, LVH: left ventricular hypetrophy, NT-proBNP: N-terminal-pro brain natriuretic peptide.

Our cellular model suggests that enzyme deficiency and accumulation of unmetabolized Gb3 in atrial iPSC-CMs are associated with key alterations in cellular contractility, intracellular calcium handling, AP morphology and DAD occurrence. Similar findings have been demonstrated in ventricular iPSC-CMs from patient-derived iPSCs with FD including higher spontaneous AP frequency, shorter AP duration, increased sodium current density, and increased upstroke velocity; all suggesting increased excitability (28). The same study demonstrated disruption to intracellular calcium handling, with transients exhibiting a greater amplitude and reduction in peak-width duration. This study, however, did not investigate atrial iPSCs and was not guided by *in-vivo* clinical data. We have demonstrated, for the first time, alterations in the atrial APs in *GLA* p. *N215S* iPSC-CMs. The shifting of diastolic membrane potential to a more positive value which is closer to the threshold potential for depolarization is pro-arrhythmic as it causes an increases myocyte excitability and may contribute to an increased risk of ectopic activity (29). We have also demonstrated an increased AP upstroke velocity in atrial *GLA* p. *N215S* iPSC-CMs compared to WT. The AP upstroke is largely controlled by voltage-gated sodium channels driving the inward sodium current (30). An increase in conduction velocity, while generally considered anti-arrhythmic, has been proposed as a substrate for arrhythmia in atrial and ventricular CMs (12). It has been suggested that rapid intra-atrial conduction velocity counteracts the fast inter-atrial conduction via Bachmann’s bundle, allowing for co-ordinated atrial depolarization. A shorter AP with a quick upstroke has also been demonstrated in patient-derived stem cell-derived ventricular CMs in FD (28). Our findings reflect this in atrial CMs and provide a mechanism for the observation of P-wave duration and PQ shortening as the earliest electrical manifestation of FD cardiomyopathy on 12-lead ECG (10). The ECG data from adults with FD together with our data from atrial iPSC-CM, suggests intrinsic early electrical changes which may be secondary to Gb3 accumulation.

Contraction analysis results demonstrate significantly higher force of contraction and peak amplitude with prolonged peak-to-peak times in *GLA* p. *N215S* iPSC-CMs compared with WT (21). Our study, for the first time, demonstrates intrinsic changes to the properties of atrial myocytes at a cellular level manifesting as increases in contraction force that are not due to passive effects of the elevated LVEDP and LVH. Published clinical data demonstrates that impairments in LA strain (contractile function) were weakly associated with parameters of LV diastolic and systolic function, implying that long standing atrial structural changes accounting for LA strain impairment in FD overcoming and negating the cellular effects which tend to increase contractile force. (6).

The changes observed in this study of a slower beat rate in *GLA* p. *N215S* iPSC-CMs compared to WT are in keeping with published literature which records that patients with FD are often bradycardic. Our data suggest the bradycardia may be a result of atrial and perhaps sinus node cellular changes caused by Gb3 accumulation.

Another novel finding of this study is the identification of alterations in intracellular calcium handling in atrial *GLA* p. *N215S* iPSC-CMs including a greater transient amplitude and prolonged duration. Although it should be noted that amplitude measurements from optical mapping of non-ratiometric indicators are influenced by several non-physiological factors, including inhomogeneous loading and excitation, these results suggest an overall increase in calcium that is cycled into and out of the cytosol with each excitation (31). This may explain the increase in contraction force observed in these cells, and also aligns with published data in patient derived ventricular iPSC-CMs in FD which demonstrate a similar finding of greater amplitude and a greater rising slope with reduced peak width duration (28). Furthermore, the finding of a greater number of atrial *GLA* p. *N215S* iPSC-CMs exhibiting DAD activity compared to WT is consistent with a greater quantity of intracellular calcium which predispose to DADs (32, 33). DADs can trigger focal activity that can lead to arrhythmogenesis, a potential mechanism explaining the increased arrhythmia-burden in these patients, including for AF (34). The higher level of DADs in *GLA* p. *N215S* iPSC-CMs may relate to a shift of the diastolic membrane potential in atrial APs to a more positive value, closer to the depolarization threshold, increasing myocyte excitability and focal ectopic activity risk. Further investigations are required to directly assess subcellular calcium dynamics in these cells.

When reproducing the observed calcium handling and action potential findings from *GLA* p. *N215S* iPSC-CMs in an *in-silico* atrial population, we found simulated P-waves with a shorter P-wave duration and PQ interval than P-waves without the cellular findings imputed. The P wave morphology was similar to that seen in the P-waves of patients with FD stage 1. This striking observation provides further evidence that intrinsic changes in atrial calcium handling and electrophysiology may, account for the abnormalities in P-wave duration and PQ interval seen in early FD cardiomyopathy. We also demonstrate an increase in the APD restitution slope, the appearance of APD alternans and (in our bi-atrial tissue model) AF inducibility by ectopic activity in the pulmonary veins. APD restitution is the rate-dependant adaptation of the AP (35). Changes in APD restitution may provoke re-entrant arrythmia particularly in the context of a prolonged repolarization phase and in conjunction with premature stimuli (36, 37). Our ECG analysis demonstrated increased PAC occurrence in FD patients likely arising from ectopic activity in the pulmonary veins. Subsequently, in our FD bi-atrial model informed by our observed clinical and cellular findings, AF was induced by two ectopic beats originating from the left superior pulmonary vein. These findings confirm the pro-arrhythmic nature of observed cellular changes induced by *GLA* p. *N215S* FD. The *in-vivo, in-vitro* and modelling experiments used in this model for FD may be applicable to other atrial cardiomyopathies with the potential of developing atrial arrhythmia-specific therapy

## LIMITATIONS

This study involves one of the largest datasets of patients with FD. However, when comparing to studies on related cardiac conditions, the cohort size is comparatively small since FD is a rare disease. The main limitation of the cellular data is that there can be several variables in the differentiation process when culturing atrial cardiomyocytes which affects the quality and subsequent molecular and functional properties of the cells. To mitigate this, we conducted experiments on multiple batches (minimum of 3) of high-quality cells in a coordinated beating monolayer, suitable for analysis.

## CONCLUSION

In summary, in this study of adults with FD and atrial iPSC-CMs with the *GLA* p. *N215S* variant for FD, we confirm early P-wave changes in “phenotype negative” patients. In an atrial myocyte model of FD we found novel cellular changes including alterations in contraction force, intracellular calcium handling, and atrial electrophysiology which may account for the early electrical changes observed on 12-lead ECGs of FD patients. In an *in-silico* atrial model, the cellular changes produced a similar P-wave morphology to that seen in clinical early-stage FD cardiomyopathy and predicted an increase in AF vulnerability. This study provides new insights into the underlying mechanisms contributing to the arrhythmic substrate in FD and validates the use of iPSC models to understand arrhythmia mechanisms. These identified changes may act as targets for cardioprotective therapy to reduce the burden of arrhythmia and stroke in FD.

## NOVELTY AND SIGNIFICANCE

### What is known?

- Arrhythmia and stroke are responsible for large burden of cardiovascular mortality in FD. Whilst predisposing factors and pathophysiological mechanisms for ventricular arrhythmia are established, there is less evidence for mechanisms of atrial myopathy and arrhythmia. Yet, AF prevalence is high and likely a significant contributor to stroke burden.
- The earliest detectable cardiac manifestations on ECG are shortening or prolongation of the PQ interval. The mechanisms of PQ shortening are poorly understood.
- *GLA* p. *N215S* is a non-classical cardiac variant of FD, previously thought to be “late onset”, with manifestations predominantly presenting late in adulthood.

### What new information does this article contribute?

- Signal-averaged P-wave analysis in adults with FD of varying stages of cardiomyopathy demonstrate P-wave duration and PQ shortening in adults with FD, but no overt cardiovascular phenotype. P-wave duration and PQ interval prolong as cardiomyopathy progresses, likely due to atrial structural remodelling including dilatation and impaired atrial strain.
- We have identified several mechanisms for atrial arrhythmogenesis in FD. We demonstrate alterations in contraction, calcium handling and excitability in gene-edited *GLA* p. *N215S* atrial IPSC-cardiomyocytes which under-express α-GAL A and accumulate Gb3, compared to WT. We confirm arrhythmia inducibility when imputing our cellular findings in an *in-silico* atrial model.
- The atrial iPSC-CM data supports the early clinical ECG findings in FD, suggesting electrical instability and remodelling in atrial cardiomyocytes begins early and progresses insidiously, forming a substrate for atrial arrythmia. The combined use of *in-vivo* and *in-vitro* data with an *in-silico* model may be applicable to other atrial cardiomyopathies.

In summary, we confirm for the first time, P-wave duration and PQ interval shortening in FD patients with a ‘pre-disease’ cardiac phenotype, compared to age-matched healthy controls. Supporting this, we confirm accumulation of un-metabolised Gb3 substrate in gene-edited *GLA* p. *N215S* atrial cardiomyocytes which are deficient in α-GAL A. We found changes in atrial electrophysiology, contraction, and calcium handling, which may be explained by early accumulation of Gb3, altering the properties of the atrial cardiomyocytes. Electrical and calcium handling alterations seen in the *N215S* atrial CMs were sufficient to increase arrhythmia risk in *in-silico* atrial models. These insights into early atrial alterations suggest a need for increased AF monitoring of patients with early cardiac stage. The mechanisms identified in this study support further research into targeted therapy to reduce the atrial substrate for arrhythmia, thus reducing the prevalence of AF and stroke in FD.

## ACKNOWLEDGMENTS

The authors would like to acknowledge the support and work of the Centre for Rare Diseases and Department of Echocardiography at the Queen Elizabeth Hospital, Birmingham, United Kingdom.

